# PlasmidEC and gplas2: An optimised short-read approach to predict and reconstruct antibiotic resistance plasmids in *Escherichia coli*

**DOI:** 10.1101/2023.08.31.555679

**Authors:** Julian A. Paganini, Jesse J. Kerkvliet, Lisa Vader, Nienke L. Plantinga, Rodrigo Meneses, Jukka Corander, Rob J.L. Willems, Sergio Arredondo-Alonso, Anita C. Schürch

## Abstract

Accurate reconstruction of *Escherichia coli* antibiotic resistance gene (ARG) plasmids from Illumina sequencing data has proven to be a challenge with current bioinformatic tools. In this work, we present an improved method to reconstruct *E. coli* plasmids using short reads. We developed plasmidEC, an ensemble classifier that identifies plasmid-derived contigs by combining the output of three different binary classification tools. We showed that plasmidEC is especially suited to classify contigs derived from ARG plasmids with a high recall of 0.941. Additionally, we optimised gplas, a graph-based tool that bins plasmid-predicted contigs into distinct plasmid predictions. Gplas2 is more effective at recovering plasmids with large sequencing coverage variations and can be combined with the output of any binary classifier. The combination of plasmidEC with gplas2 showed a high completeness (median=0.818) and F1-score (median=0.812) when reconstructing ARG plasmids and exceeded the binning capacity of the reference-based method MOB-suite. In the absence of long read data, our method offers an excellent alternative to reconstruct ARG plasmids in *E. coli*.

**Data Summary:** No new sequencing data have been generated in this study. All genomes used in this research are publicly available at the GenBank and Sequence Read Archive of the National Center for Biotechnology Information. Accession numbers are specified in Supplementary Materials.

Scripts to reproduce the results reported in this manuscript can be accessed at https://gitlab.com/jpaganini/ecoli-binary-classifier. The ensemble classifier, plasmidEC, is publicly available at https://gitlab.com/mmb-umcu/plasmidEC (release 1.3.1), and gplas2 (release 1.0.0) can be found at https://gitlab.com/mmb-umcu/gplas2.

**Impact Statement:** *Escherichia coli* has emerged as a highly pervasive multidrug resistant pathogen on a global scale. The dissemination of resistance is significantly influenced by plasmids, mobile genetic elements that facilitate the transfer of antimicrobial resistance genes within and between diverse bacterial species. Consequently, precise and high-throughput identification of plasmids is imperative for effective genomic surveillance of resistance. However, accurate plasmid reconstruction remains challenging with the use of affordable short-read sequencing data. In this work, we present a novel method to accurately predict and reconstruct *E. coli* plasmids based on Illumina data. Additionally, we demonstrate that our approach outperforms the reference-based method MOB-suite, especially when reconstructing plasmids carrying antimicrobial resistance genes.

## Introduction

*Escherichia coli* is a commensal gram-negative bacterium inhabiting the gastrointestinal tract but is also the leading cause of bloodstream and urinary tract infections in humans [1,2]. In recent years, the emergence and spread of multidrug resistant *E. coli* lineages limits the treatment options for such infections [3,4]. Moreover, a recent assessment of the global burden of antimicrobial resistance (AMR) estimated that AMR *E. coli* infections accounted for more than 250,000 deaths in 2019, placing *E. coli* as one of the most prevalent AMR pathogens worldwide [5].

Horizontal gene transfer is one of the main drivers behind the rapid spread of AMR [6–8]. Antibiotic resistance genes (ARGs) are commonly associated with mobile genetic elements (MGEs), which facilitate their mobility across bacteria [9,10]. Out of these MGEs, plasmids play a pivotal role by disseminating AMR in clinical settings as well as in other environments [11–13]. Plasmids are frequently transmitted among bacteria of the same species, but they can also be shared between bacteria of different species or even different genera [14–17]. Given their relevance in the spread of AMR genes, it is critical to develop high-throughput methods to identify plasmids in a precise, fast and accessible manner.

Bacterial genomes have been massively studied using short-read sequencing platforms. However plasmids tend to contain repetitive elements that cannot be spanned by short-reads and thus their sequence is usually fragmented into several contigs and mingled with other genomic elements. This makes it hard to reconstruct complete plasmids from short-read sequencing data [18].

Several fully-automated bioinformatic tools are currently available to predict plasmids from short-read sequencing data. They can be broadly categorised into two groups: (i) tools that produce a binary classification of contigs as either plasmid- or chromosome-derived, predicting the total plasmid content of a bacterial strain, often referred to as the ‘plasmidome’ (without reconstructing individual plasmids), and (ii) tools that aim to recover complete sequences for individual plasmids [19]. The latter group, termed plasmid reconstruction tools, provides a more suitable output for plasmid epidemiology studies.

We recently evaluated the performance of several plasmid reconstruction tools for use with *E. coli* short-read data [19]. We found that the best performing tool, MOB-suite [20], only achieved the correct reconstruction of 50.2% of the plasmids. Moreover, all tools underperformed when attempting to reconstruct plasmids containing antibiotic resistance genes (ARG-plasmids), ranging from 3.4% to 27.9% correct ARG-plasmid reconstructions. These results emphasised the need to improve current methods to predict ARG-plasmids in *E. coli*.

Here, we present a new high-throughput method to reconstruct *E. coli* plasmids from short-read sequencing data. Firstly, we optimised gplas [23], a plasmid binning tool, to compute walks in the assembly graph corresponding to plasmids with a pronounced coverage variation. Secondly, we developed an ensemble classifier, plasmidEC, combining multiple existing binary classification tools (Plascope [21], RFplasmid [22], Platon [23] and mlplasmids [24]) to predict plasmid-derived contigs. Coupling plasmidEC with gplas2 allowed to accurately bin plasmid-derived contigs into separate components corresponding to individual plasmid sequences. Our method outperforms all currently available plasmid reconstruction tools for *E. coli*, especially for predicting ARG-plasmids.

## Methods

All scripts used to reproduce the analyses can be found at gitlab.com/jpaganini/ecoli-binary-classifier. R version 3.6.1. was used for all R scripts.

### Benchmark datasets

A dataset of 240 complete *E. coli* genomes from 8 different phylogroups and 117 sequence types (STs), carrying 631 plasmids, was selected as previously described in Paganini et al. [19]. Samples were isolated from animals, humans and the environment, resulting in a diverse dataset with respect to phylogeny and plasmid content. All genome sequences were completed by the combination of short- and long-read sequencing data. Short-read sequences and complete genomes were downloaded from NCBI using SRA tools (v2.10.9) and ncbi-genome-download (v0.2.10) (https://github.com/kblin/ncbi-genome-download), respectively. Genomes present in the training datasets or reference databases of existing plasmid classification tools (mlplasmids, PlaScope, Platon and/or RFPlasmid) were removed (n=26). The remaining 214 samples, carrying 542 plasmids, were used to benchmark the binary classifiers (Supplementary Data 1). From these, 15 genomes (Supplementary Data 2) were randomly selected for optimisation of the gplas algorithm and excluded from later comparisons. The remaining genomes (n=199, 483 plasmids) were used to benchmark the plasmid reconstruction methods.

#### Benchmarking binary classification tools and construction of plasmidEC

##### Selection of contigs for benchmarking

Short-read sequences of each sample were assembled with bactofidia (v1.1) (https://gitlab.com/aschuerch/bactofidia), a pipeline that relies on SPAdes for genome assembly (v3.11.1)[25]. The resulting contigs (n=18,963) were labelled as chromosome- or plasmid-derived by alignment to their respective complete genomes using QUAST (v5.0.2)[26]. Only contigs larger than 1,000 bp with an alignment of at least 90% the contig length were considered (n=15,020). Of those, contigs aligning to multiple positions in the genome (ambiguously aligned contigs) were included as long as they exclusively aligned to either the chromosome or to plasmids (n=1,236). The same criterion was used for the inclusion of misassembled contigs (n=1,862). In total, the benchmark dataset included 14,746 contigs (Supplementary Figure S1).

##### Assessment of the individual binary classifiers

Contigs were classified by mlplasmids (v2.1.20), PlaScope (v.1.3.121), Platon (v.1.619) and RFPlasmid (v.0.0.1722). All tools were run using default parameters. We assessed the performance of the four binary classifiers by comparing, for each contig, their prediction to the true class of the contig, as described in the section above. For PlaScope, an ‘unclassified’ prediction was handled as a negative prediction. Predictions were categorised into: True Positives (TP, prediction = plasmid, class = plasmid), True Negatives (TN, prediction = chromosome, class = chromosome), False Positives (FP, prediction = plasmid, class = chromosome) and False Negatives (FN, prediction = chromosome, class = plasmid). Global performance of the tools was evaluated with the following metrics:

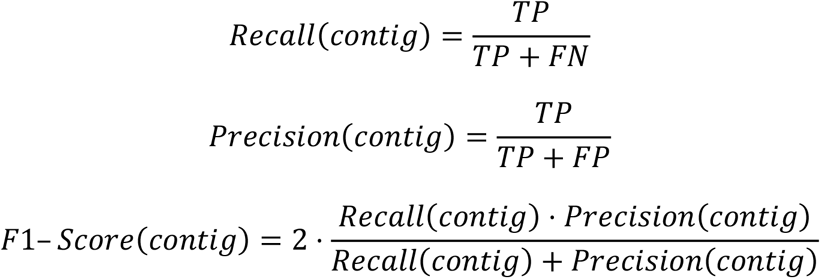

##### Assessment of the ensemble classifiers

To improve the predictions obtained by independent tools, we combined their output into distinct ensemble classifiers that implemented a majority voting system. We tested four different combinations of individual classifiers: mlplasmids/PlaScope/Platon, mlplasmids/PlaScope/RFPlasmid, mlplasmids/Platon/RFPlasmid and PlaScope/Platon/RFPlasmid. A final classification of each contig (chromosome or plasmids) was obtained by combining the output of the tools using an R script (provided in the accompanying code repository). The ensemble classifiers were evaluated using the same metrics as described above.

##### Construction of plasmidEC

The tool consists of a bash wrapper script that automatically installs and runs all required individual classifiers and combines their results with a majority voting system. Based on the performance for *E. coli*, the combination of PlaScope/Platon/RFPlasmid was selected as the default. PlasmidEC is publicly available at https://gitlab.com/mmb-umcu/plasmidEC.

#### Benchmarking plasmid reconstruction tools

##### Running plasmid predictions tools

Prior to assembly, Illumina raw reads were trimmed using trim-galore (v0.6.6) (https://github.com/FelixKrueger/TrimGalore) to remove bases with a Phred quality score below 20. Unicycler (v0.4.8) [27] was then applied to perform *de novo* assembly with default parameters. Contigs larger than 1,000 bp were used as input for MOB-suite (v3.0.0) [20], while assembly graphs in GFA format served as input for gplas2 (v2.0.0). To run gplas2, nodes from the graph were first classified as plasmid- or chromosome-derived using either plasmidEC or PlaScope; only nodes larger than 1,000 bp were classified. Output from the tools was modified to assign probabilities for the classification of each node, which is required by the gplas algorithm. For PlaScope, discrete probabilities were assigned based on the node classification status; if a node was classified as plasmid, a probability of 1 was assigned, while chromosome-predicted nodes were assigned zero. In the case of unclassified nodes, a probability of 0.5 was assigned. By default, plasmidEC assigns probabilities based on the fraction of tools that agreed on the classification. For example, if two out of three tools agreed in classifying a node as plasmid, a probability of 0.66 is assigned.

##### Analysis of the plasmid bin composition

To evaluate the bins created by MOB-suite and gplas2, we used QUAST (v5.0.2) [26] to align the contigs of each bin to the respective complete reference genome. We calculated accuracy, completeness and F1-score on the base-pair level, as specified below.

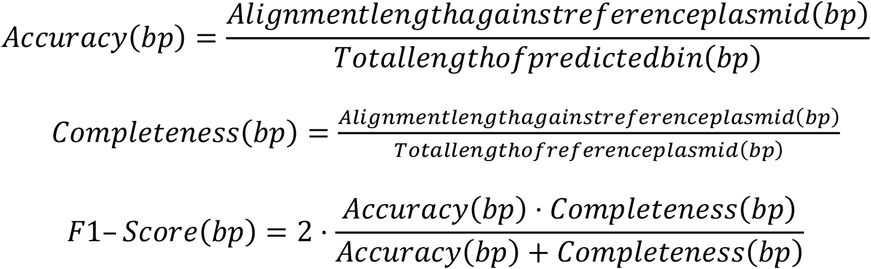

If a bin was composed of contigs derived from different plasmids, then accuracy_(bp)_, completeness_(bp)_ and F1-score_(bp)_ were reported for each plasmid-bin combination.

We also evaluated the number of reference plasmids that were detected by each tool. We consider a reference plasmid as detected when at least a single contig of the plasmid was included into the predictions.

To determine *combined completeness* for each reference plasmid, all bins generated in an isolate were combined as follows:

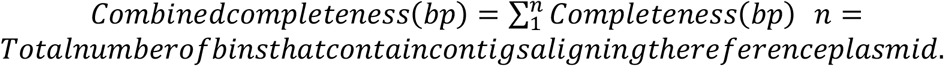

##### Antibiotic Resistance Gene (ARG) Prediction

Resistance genes were predicted by running Abricate (v1.0.1) against the Resfinder [28] database (database indexed on 19 April 2020) with reference plasmids as query, using 80% as identity and coverage cut-off. The same software and parameters were used to predict the presence of ARGs in the plasmid-predicted contigs bins generated by each of the plasmid reconstruction tools.

##### Evaluation of ARGs binning

For bins that carried ARGs, we calculated Recall(ARG) and Precision(ARG) as indicated below.

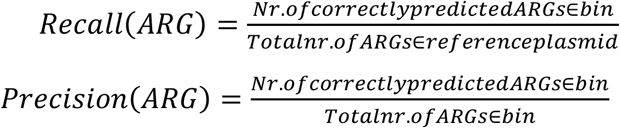

##### Evaluating unbinned nodes in gplas predictions

Unitigs classified as unbinned by gplas (n=78) were aligned to the corresponding complete reference genome using QUAST (v5.0.2). The results of these alignments were used to determine the origin of the unitig (plasmid or chromosome). For isolates that contained more than one unbinned unitig (n=19), coverage information of all unitigs (bin and unbinned) was extracted from the header of the FASTA files generated after unicycler assembly. From these data, coverage variance for all replicons was calculated and plotted using R (v.3.6.1).

##### Evaluating the recovered fraction for each reference plasmid

We calculated the maximum completeness(bp) that can be obtained to reconstruct every reference plasmid using short-read sequencing data. Before applying any classification tool, all nodes from the assembly graph were converted to FASTA format using the ‘extract’ option of gplas2. Nodes smaller than 1,000 bp or smaller than 500 bp were filtered out using seqtk (v1.3) (https://github.com/lh3/seqtk), and remaining nodes were aligned to their respective complete reference genomes using QUAST to obtain the completeness(bp) values. The completeness(bp) value was called the *recovered fraction*.

##### Read coverage of missing reference plasmids

A small number of plasmids were either completely missed or recovered with low completeness after short-read assembly. In order to determine if these sequences were also missing from short-reads, trimmed Illumina reads were aligned to reference genomes using BWA MEM (v.0.7.17) [29] with default parameters. Resulting SAM files were converted to BAM and sorted using SAMtools (v1.9) [30]. Read coverage per base was determined using BEDTOOLS (v2.30.0) [31].

## Results

### Optimisation of gplas to improve the reconstruction of *E. coli* plasmids

Gplas is an algorithm that performs *de novo* reconstruction of plasmids through multiple steps (Figure 1 - Steps 1 to 3) [32]. In short, nodes from the assembly graph are initially classified as plasmid-derived or chromosome-derived by an external binary classification software, which also assigns a probability to the classifications. Then, plasmid-predicted unitigs act as seeds to compute plasmid walks with homogeneous coverage in the assembly graph using a greedy approach. Finally, these unitigs are binned together into individual components based on their co-existence in the computed plasmid walks. A detailed description of the algorithm can be found in the original publication [32]. Given that gplas performed sub-optimally when reconstructing *E. coli* plasmids in our previous study [19], in gplas2 we introduced two major modifications to the algorithm:

**Figure 1.**
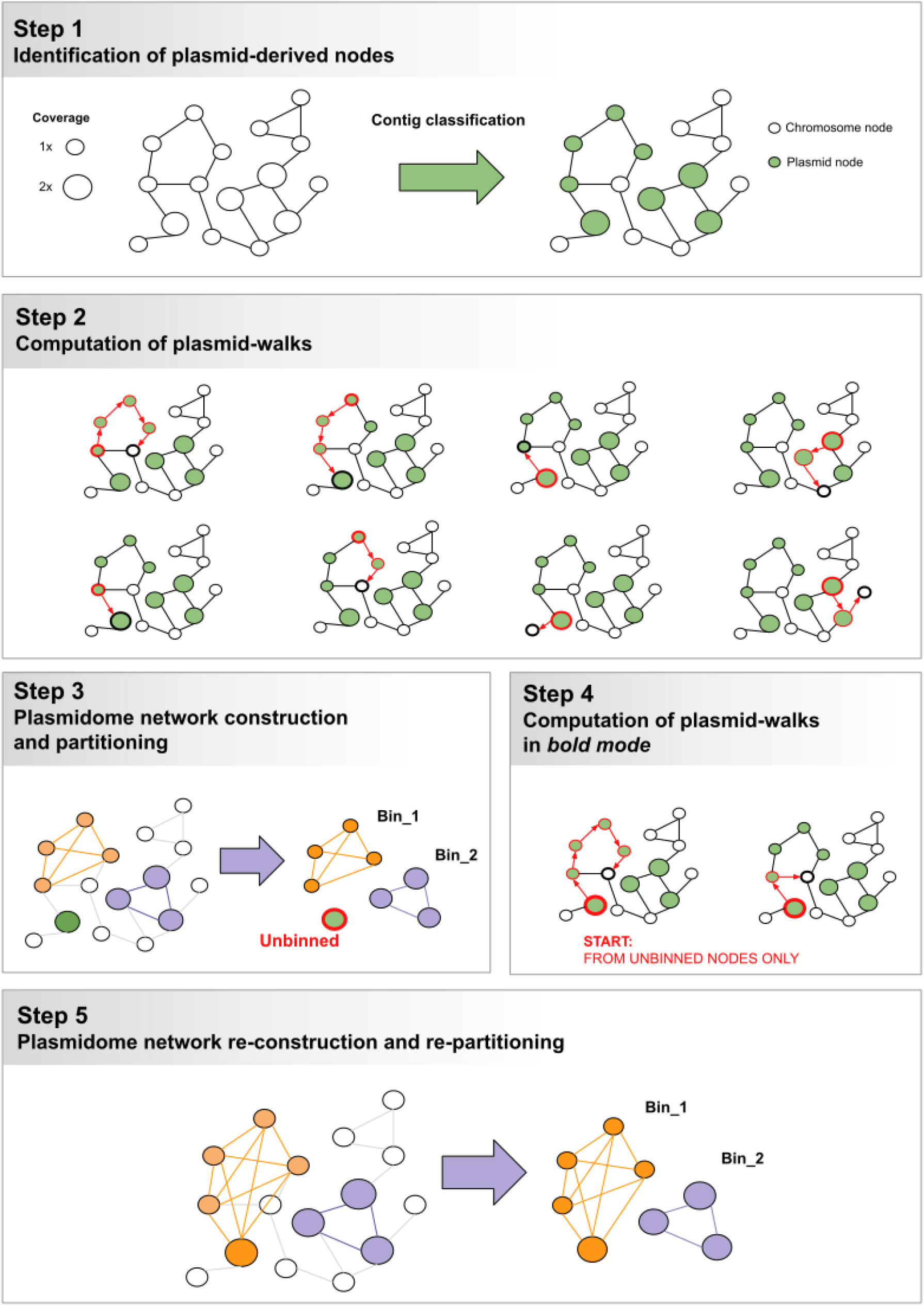
Schematics on gplas2 algorithm. The steps 4 and 5 were added to gplas2 in order to recover unbinned unitigs.

#### A) Expansion of the input options for binary classification

Coupling gplas with an accurate binary classifier improves the reconstruction of plasmids, as we previously demonstrated for *Enterococcus faecalis* and *Klebsiella pneumoniae* [32,33]. Consequently, the gplas2 algorithm accepts predictions from any binary classifier, provided they output classification probabilities and expected file formats.

#### B) Re-iterating plasmid walks over initially unbinned contigs

Gplas constructs plasmid walks over the assembly graph to connect unitigs that potentially originate from the same plasmid (Figure 1 - Step 2). Consequently, plasmid-predicted unitigs that can’t be connected to other unitigs through these walks are classified as unbinned, and are not included in the plasmid predictions (Figure 1 - Step 3). Unbinned unitigs seem to originate from reference plasmids that were sequenced with a pronounced coverage variation (Supplementary Figure S2). This sequencing artefact poses a challenge to the gplas algorithm, which builds plasmid walks from unitigs with homogeneous coverage. Consequently, we modified gplas to consider these coverage variations (Figure 1 - Steps 4 & 5). Whenever unbinned unitigs are produced, gplas2 will generate a second round of binning in bold mode by running two additional steps:

##### 1) Computation of plasmid walks in bold mode starting from unbinned unitigs

If unbinned unitigs are predicted, new bold plasmid walks will be constructed. When creating the bold walks, a higher coverage variance threshold between plasmid-predicted unitigs is allowed. This threshold can be defined by the user and is a multiple of the coverage variance observed for chromosome-predicted unitigs. Only bold plasmid walks that start from unbinned unitigs will be retained to use in the next step, while the rest will be discarded (Figure 1 - Step 4).

##### 2) Plasmidome network reconstruction and repartitioning

Plasmid walks produced during bold mode are merged with plasmid walks from normal mode. Based on these combined data, plasmidome networks are reconstructed and repartitioned (Figure 1 - Step 5) to create new bins, using the same algorithms as in step 3.

We optimised the predictions obtained with gplas2 using a subset of 15 *E. coli* genomes that contained unbinned unitigs and that were excluded from subsequent benchmarking efforts (Supplementary Data 2). For bold walks, we allowed a coverage variance of 5, 10, 15 or 20 times the coverage variance observed for the chromosome-predicted unitigs. Plasmid predictions made with gplas2 exhibited consistently higher completeness(bp) values when compared to the original predictions (Supplementary Figure S3 A). Surprisingly, altering the coverage variance threshold above 5 did not impact completeness(bp) values. In contrast, accuracy(bp) values decreased when allowing a higher coverage variance. The highest F1-Score(bp) values (median=0.78, IQR=0.47 - 0.96) were obtained when using a coverage variance threshold of 5. Consequently, 5 was defined as the default value to construct bold plasmid walks. As a single example, we display the plasmid predictions obtained with and without running bold mode for genome GCA_013823335.1_ASM1382333v1 (Supplementary Figure S3 B and S3 C). In this case, the bold walks allowed to recover 7 additional contigs belonging to plasmids CP057179.1 and CP057180.1.

Gplas2, including the aforementioned features and a detailed user guide, can be found at https://gitlab.com/mmb-umcu/gplas2.

### Comparing binary classification methods for *E. coli*

In order to combine gplas2 with the best available binary classifier for *E. coli*, we compared the performance of four different tools (PlaScope, RFPlasmid, mlplasmids and Platon). The benchmark dataset consisted of 14,746 contigs. Of these contigs, 87.3% (n=12,872) were chromosome-derived and 12.7% (n=1,874) were plasmid-derived, as determined by alignment to complete reference genomes.

We evaluated the number of contigs which were correctly and incorrectly classified by each of the tools and calculated recall_(contig)_, precision_(contig)_ and F1-score_(contig)_ (Supplementary Table S1). Plascope was able to correctly identify the highest number of plasmid-derived contigs (True Positives, n=1,629), while the rest of the tools detected between 1,297 and 1,523 plasmid-derived contigs. Notably, PlaScope also included the least chromosomal contamination in its predictions (False Positives, n=117), closely followed by Platon (n=122). In contrast, mlplasmids and RFPlasmid included a higher amount of chromosome-derived contigs in their plasmidome predictions (n=418 and n=420, respectively). PlaScope was the tool with the highest F1-score_(contig)_ (0.900) followed by Platon (0.861), RFPlasmids (0.798) and mlplasmids (0.722). For most tools, precision_(contig)_ values were higher than recall_(contig)_ values, indicating that the predicted plasmidome mostly consists of true plasmid-derived contigs, but also that plasmid contigs were frequently missed by the tools.

We also explored the congruence in contig classifications across tools (Figure 2). All tools agreed on the correct classification of 51.8% of plasmid-derived contigs (True Positives: n=971, Figure 2A), and another 26.5% plasmid-derived contigs were correctly classified by at least three tools (n=497). Also, a high fraction (94.1%) of chromosome-derived contigs were correctly classified by all tools (True Negatives: n=12,116, Figure 2B). Moreover, only a minority of plasmid-derived and chromosome-derived contigs were missed by most of the tools and correctly classified by just a single tool (True Positives: 85/1,874, 4.7%, True Negatives: 58/12,872, 0.5% respectively). From these observations, we concluded that contig misclassifications are primarily derived from individual tools (Figure 2C and 2D).

**Figure 2.**
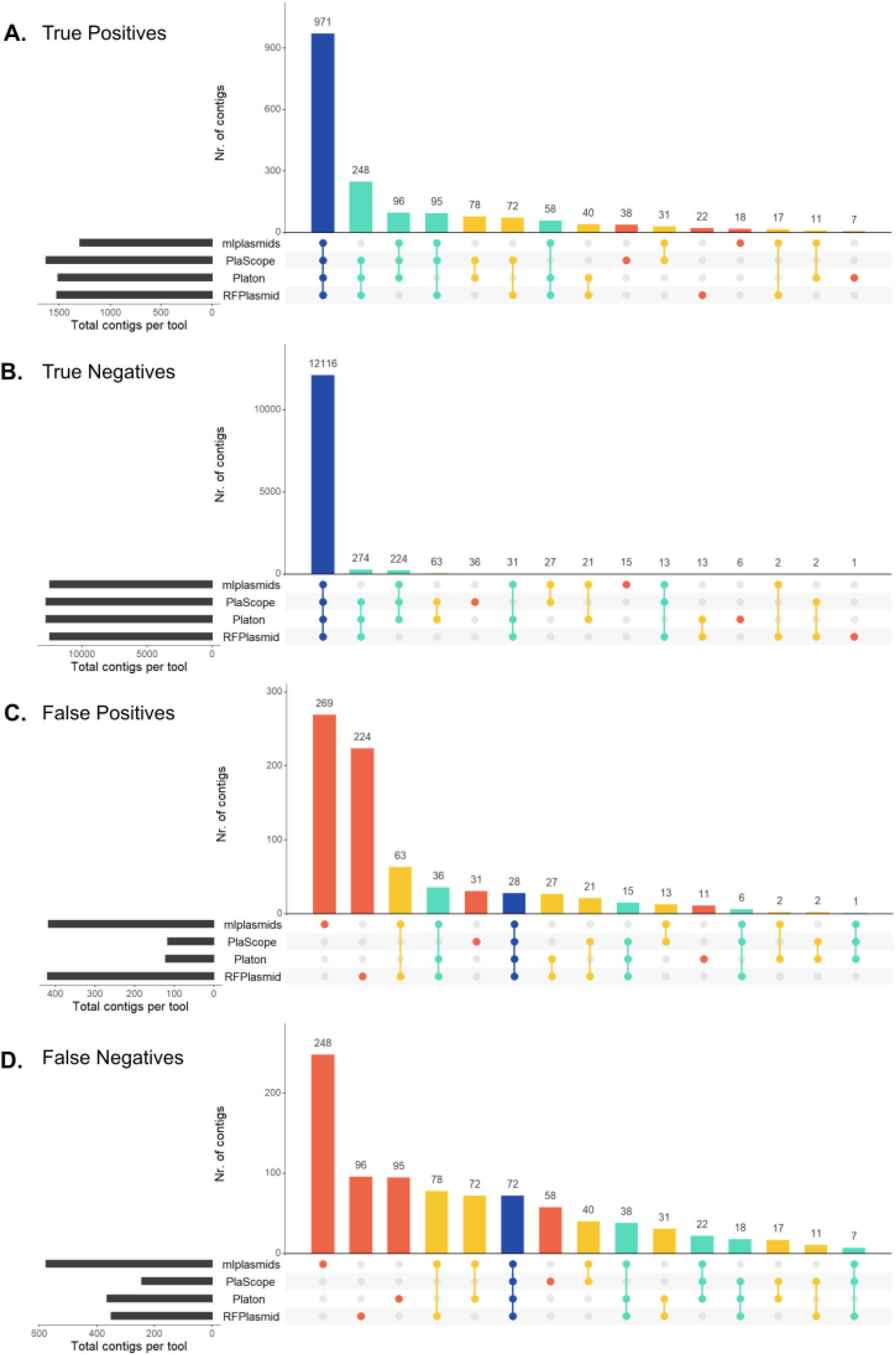
Upset diagrams showing congruence in contig classification by different binary prediction tools (absolute counts). True Positives (TP; prediction=plasmid, class=plasmid), True Negatives (TN; prediction = chromosome, class=chromosome), False Positives (FP; prediction=plasmid, class=chromosome), False Negatives (FN, prediction=chromosome, class=plasmid). Bar colours indicate the number of tools that concur in the classification of the contigs.

### PlasmidEC: A voting classifier for improved detection of ARG-plasmid contigs in *E. coli*

We theorised that discarding software-specific misclassifications, while keeping correct classifications shared by multiple tools, could improve the overall binary classification of *E. coli* contigs as plasmid- or chromosome-derived. To explore this, we combined the predictions of three individual classifiers and extracted their majority vote as the final classification.

After testing all possible combinations of individual classifiers, we found that Platon/PlaScope/RFPlasmid displayed the highest overall performance of voting classifiers with the highest F1-score_(contig)_ (0.904). This ensemble classifier achieved an F1-score_(contig)_ similar to PlaScope (0.900) but had a slightly higher recall_(contig)_ (0.884 and 0.869, respectively) (Figure 3 A and B, Supplementary Table S1).

**Figure 3.**
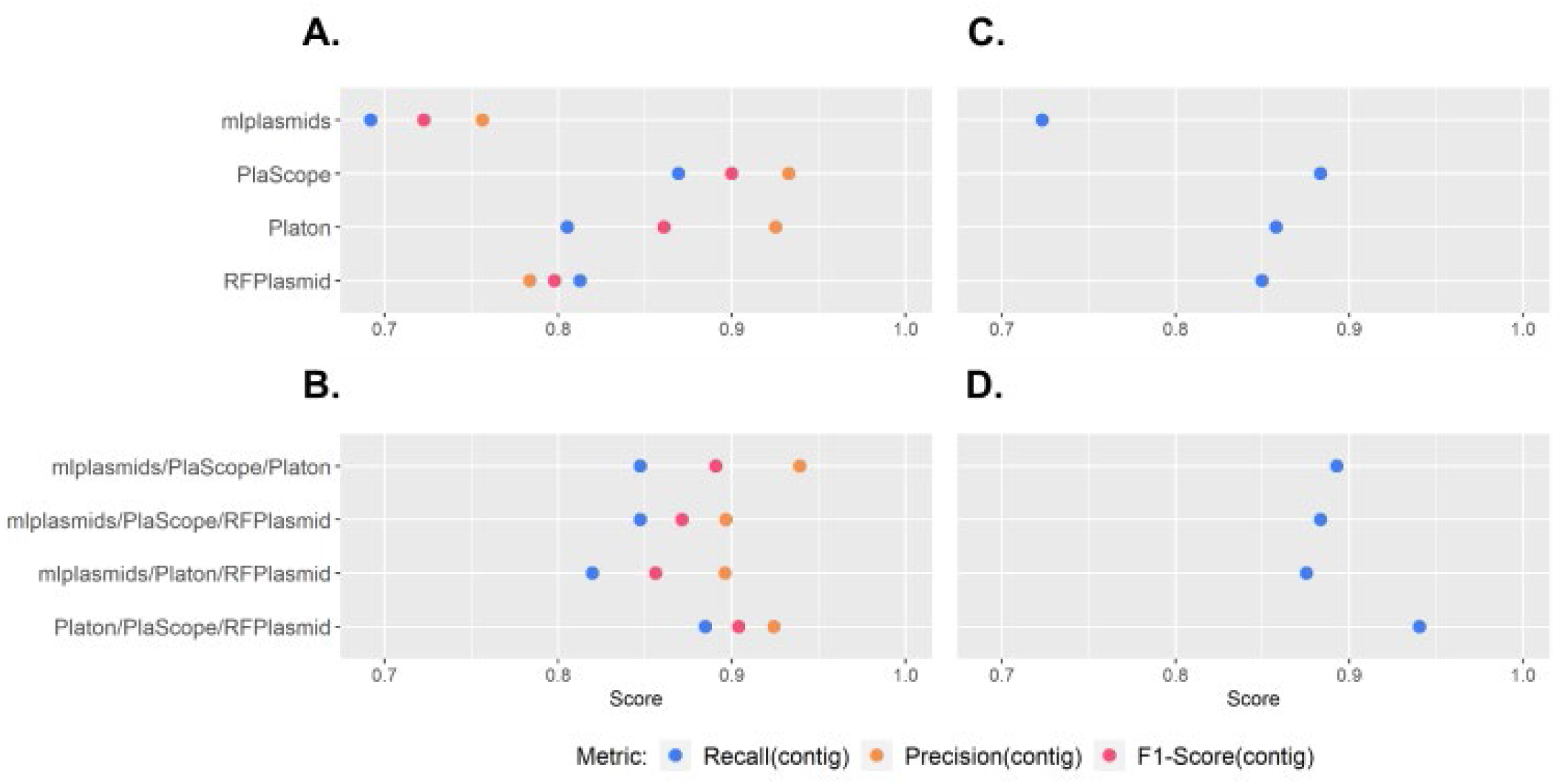
Performance of individual binary classifiers and plasmidEC combinations, measured by recall(contig), precision(contig) and F1-score(contig) A) Individual classifiers evaluated using full dataset (n=214 genomes). B) PlasmidEC combinations evaluated using full dataset C) Individual classifiers evaluated using a dataset of ARG-plasmids (n=114 plasmids). D) PlasmidEC combinations evaluated using a dataset of ARG-plasmids.

Next, we evaluated recall_(contig)_ values for a subset of plasmids (n=114) encoding antibiotic resistance genes (ARG-plasmids) (Figure 3C and 3D, Supplementary Table S2). This dataset consisted of 860 plasmid-derived contigs, derived from 91 *E. coli* genomes. The recall_(contig)_ of individual tools ranged from 0.723 (mlplasmids) to 0.884 (PlaScope), whereas the different combinations of tools in a voting classifier reached recall_(contig)_ values ranging from 0.883 (mlplasmids/Platon/RFPlasmid) to 0.941 (Platon/PlaScope/RFPlasmid).

Based on these results, the combination of Platon/PlaScople/RFPlasmid was selected as the ensemble classifier to be implemented in a novel tool termed plasmidEC, which is publicly available at https://gitlab.com/mmb-umcu/plasmidEC.

We measured the computational resources used by the ensemble and individual classifiers (Supplementary Figure S4). Binary classifiers showed considerable differences in both CPU time and memory usage. The average CPU time required per sample was lowest for PlaScope (0.2 mins) and highest for Platon (14.9 mins). Platon also used the largest amount of memory per sample (20.6 Mb). The least amount of memory was required by mlplasmids (2.7 Mb). Because plasmidEC includes the execution of three binary classifiers, time and memory requirements were high, especially when Platon was run. The combination of mlplasmids/PlaScope/RFPlasmid required the least number of resources (CPU time = 4.5 mins, memory = 9.0 Mb) and PlaScope/Platon/RFPlasmid the most (CPU time = 21.5 mins, memory = 21.4 Mb).

### Exploiting the information from the assembly graph improves correct binning of ARG plasmids

To reconstruct individual *E. coli* plasmids, gplas2 was combined with plasmidEC and PlaScope, and performance was compared against MOB-suite, which was the best-performing plasmid reconstruction tool for *E. coli* in our recent benchmark study [19]. To retain comparability with the aforementioned study, we started with the same dataset and removed 26 genomes that were present in the PlaScope database and 15 genomes that were used to improve the gplas2 algorithm. Consequently, our benchmark dataset consisted of 199 complete *E. coli* genomes, which carried 483 plasmids. A total of 213 (44.1%) plasmids were classified as small plasmids (smaller than 18,000 bp), while the remaining 270 (55.9%) were large plasmids [19]. Given our interest in predicting ARG-plasmids, and the fact that most ARGs are encoded on large plasmids (n=382/387, 98.7%), we analysed performance separately for large ARG-plasmids (n=96) and large non-ARG-plasmids (n=174).

When evaluating the reconstruction of ARG-plasmids, we found that the F1-Score_(bp)_ values of gplas2 combined with either plasmidEC (gplas2_plasmidEC) or PlaScope (gplas2_PlaScope) were similar (Figure 4A, Table 1). However, gplas2_plasmidEC (median=0.81, IQR=0.53 - 0.93) performed slightly better than gplas2_PlaScope (median=0.76, IQR=0.52 - 0.94). Notably, both gplas2 methods outperformed MOB-suite, which presented a lower F1-Score_(bp)_ (median= 0.44, IQR= 0.18 - 0.87). As accuracy_(bp)_ values were nearly identical across tools, the disparity in F1-Scores_(bp)_ can be explained due to the differences in completeness_(bp)_. In contrast, combined completeness_(bp)_ distributions were virtually identical among tools. These results suggested that all methods had a similar capacity to detect contigs derived from ARG-plasmids, but gplas2 performed better at binning these contigs together into individual predictions. This hypothesis was confirmed by analysing the number of bins into which each reference plasmids was fragmented (Figure 4B). For ARG plasmids, we found that MOB-suite fragmented 49% of plasmids into multiple predictions, while both gplas2 methods did so in only 14% of the cases.

**Table 1.** Performance summary of three plasmid prediction tools, for the prediction of different plasmid types.

|  | **MOB-suite** | **gplas2_plasmidEC** | **gplas2_PlaScope** |
| --- | --- | --- | --- |
| **Large Plasmids (n=270)** |  |  |  |
| Nr. of detected plasmids* | 263 (97.4%) | 253 (93.7%) | 254 (94.1%) |
| **ARG-Plasmids (n=96)** |  |  |  |
| F1-Score(bp) (median, IQR) | 0.421 (0.172 - 0.860) | 0.812 (0.529 - 0.934) | 0.758 (0.520 - 0.936) |
| Completeness(bp) (median, IQR) | 0.317 (0.114- 0.803) | 0.818 (0.520 - 0.924) | 0.818 (0.531 - 0.924) |
| Accuracy(bp) (median, IQR) | 0.883 (0.591 - 0.982) | 0.979 (0.564 - 1) | 0.979 (0.520 - 1) |
| Nr. plasmid-borne ARGs detected | 331 (86.6%) | 331 (86.6%) | 327 (85.6%) |
| Nr. chromosome-derived ARGs | 64 | 75 | 75 |
| Recall (ARG) (median, IQR) | 1 (0.42- 1) | 1 (0.86- 1) | 1 (0.86- 1) |
| Precision (ARG) (median, IQR) | 1 (0.82 - 1) | 1 (0.75 - 1) | 1 (0.77 - 1) |
| **Non-ARG-Plasmids (n=174)** |  |  |  |
| F1-Score(bp) (median, IQR) | 0.910 (0.378 - 0.977) | 0.921 (0.596 - 0.983) | 0.912 (0.571 - 0.983) |
| Completeness(bp) (median, IQR) | 0.879 (0.245 - 0.967) | 0.915 (0.618 - 0.972) | 0.918 (0.614 - 0.972) |
| Accuracy(bp) (median, IQR) | 0.978 (0.904 - 1) | 1 (0.958 - 1) | 1 (0.796- 1) |
| **Small Plasmids (n=213)** |  |  |  |
| Nr. of detected plasmids* | 174 (81.8%) | 184 (86.4%) | 196 (92.0%) |
| F1-Score(bp) (median, IQR) | 1 (0.985 - 1) | 1 (0.991 - 1) | 1 (0.990 - 1) |
| Completeness(bp) (median, IQR) | 1 (0.976 - 1) | 1 (0.996 - 1) | 1 (0.990 - 1) |
| Accuracy(bp) (median, IQR) | 1 (1- 1) | 1 (1- 1) | 1 (1- 1) |
| Nr. plasmid-borne ARGs detected | 5 (100%) | 5 (100%) | 5 (100%) |

\*A plasmid is considered detected if at least 1 contig is included in the plasmid predictions

**Figure 4.**
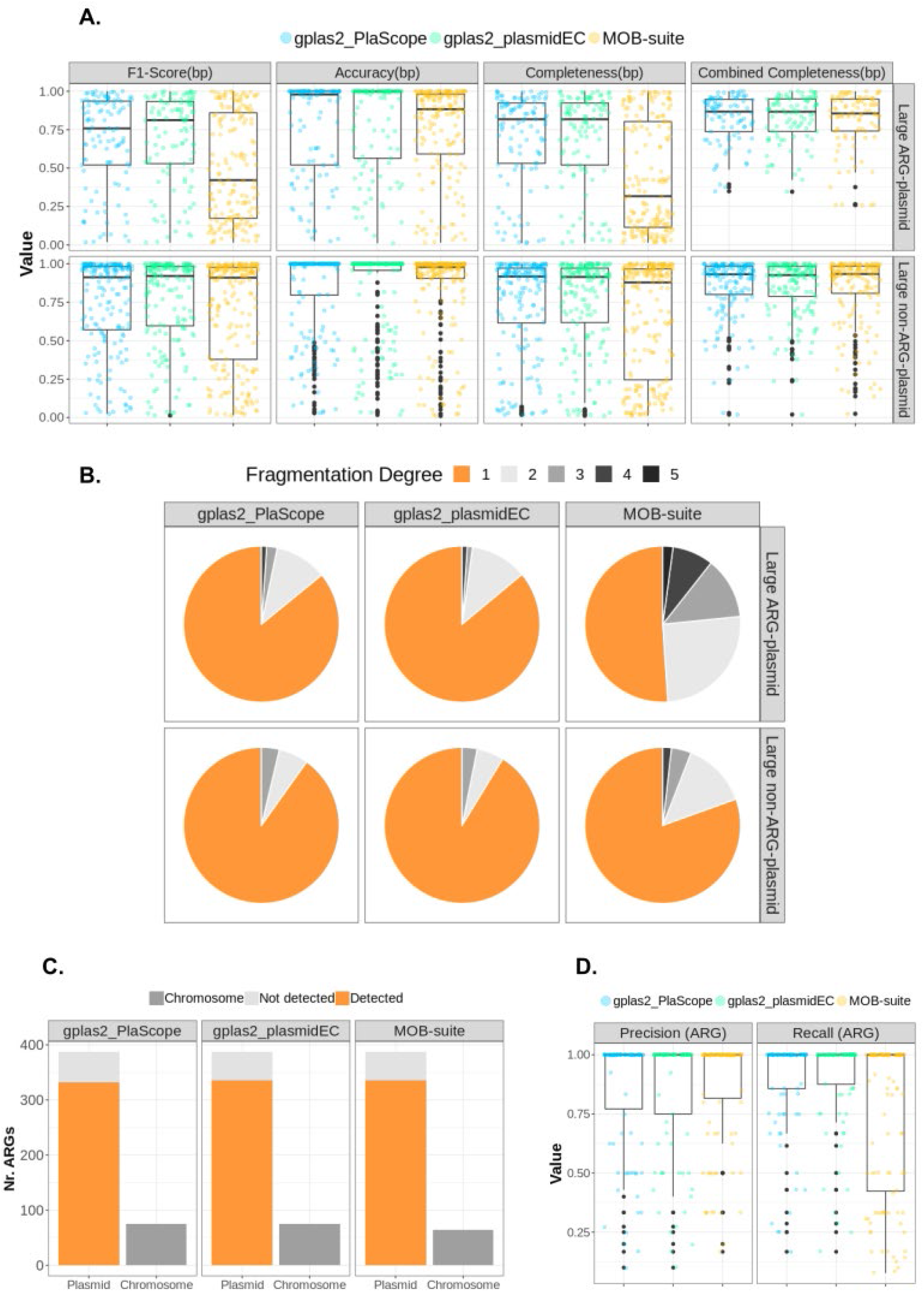
Benchmarking of plasmid reconstruction methods. A) Recall(bp), Precision(bp), F1-score(bp) and Combined Recall(bp) values for predictions corresponding to large ARG-plasmids (n=96) and large non-ARG-plasmids (n=174). B) Percentage of reference plasmids that were recovered with different fragmentation degrees (i.e. If contigs belonging to a reference plasmid are assigned to three different predictions, then the fragmentation degree equals three). C) Absolute count of ARGs included (detected) in plasmid predictions, missing ARGs (not detected) and chromosome-derived ARGs incorrectly included (Chromosome). D) Recall(ARG) and Precision(ARG) value.

All tools identified a similar number of plasmid-derived ARGs (Figure 4C). MOB-suite and gplas2_plasmidEC detected 331 (86.6%) ARGs and gplas2_PlaScope 327 (85.6%). Moreover, all tools successfully detected all ARGs present in small plasmids (n=5, 100%). In concordance with previous results, recall_(ARG)_ values (Figure 4D) for gplas2 predictions were higher than those obtained with MOB-suite (Table 1). This indicates that gplas2 performs better at correctly binning ARGs together into the same bin. However, plasmid predictions made with gplas2 also included a higher number of chromosome-derived ARGs (Figure 4C, Table 1).

Interestingly, tools performed similarly well when evaluating the reconstruction of extended spectrum beta-lactamase (ESBL) plasmids (n=42). MOB-suite reconstructions were characterised by having higher accuracy_(bp)_ and gplas2 methods reconstructed ESBL-plasmids with higher completeness_(bp)_ (Supplementary Figure S5A). Despite these differences, all tools exhibited similar F1-Score_(bp)_ values. Additionally, the number of plasmid-borne ESBL genes detected were almost identical across tools (Supplementary Figure S5B). Nevertheless, gplas2 methods performed slightly better at binning ARGs into the same prediction (Supplementary Figure S5C).

For small plasmids (n=213), all tools displayed similar performance across the three metrics, obtaining near-perfect reconstructions in all cases, with F1-score_(bp)_ medians of 1 (Supplementary Figure S6A, Table 1). This is likely due to most small plasmids being assembled into a single contig (n=196, 92.0%) (Supplementary Figure S6B), and consequently the identification of these contigs as plasmid-derived generally leads to obtaining high values for all metrics. We therefore evaluated the number of small (and large) plasmids detected by each of the tools (Supplementary Figure S6C, Table 1). Interestingly, gplas2_PlaScope detected 196 (92.0%) small plasmids, and gplas2_plasmidEC performed similarly, detecting 184 (86.4%). Both gplas2-methods outperformed MOB-suite, which detected 174 (81.79%) small plasmids.

Finally, we tested the effect of using different contig size cut-offs for plasmid reconstruction. We found no significant differences in performance of the tools when using 500 bp or 1,000 bp as the minimum contig size. A more detailed description of the results from this analysis can be found in the Supplementary Materials and in Supplementary Figures S7 - S10.

## Discussion

Accurately reconstructing *E. coli* plasmids from Illumina reads has proven to be a challenge, especially in the context of ARG-plasmids. In this work, we developed a new high-throughput method to reconstruct *E. coli* plasmids *de novo* from short-read sequencing data. Our method relies on an accurate identification of plasmid-derived nodes in the assembly graph, followed by the binning of these nodes using sequencing coverage and node connectivity information. We proved that our method outperforms other plasmid prediction tools available for *E. coli*, especially when reconstructing ARG-plasmids.

To improve the identification of plasmid-derived contigs, we built plasmidEC, an ensemble classifier that combines predictions from three individual binary classifiers and implements a majority voting system. Voting classifiers have been successfully applied in other fields of biology [35–38], but so far not for the problem of plasmidome identification. PlasmidEC correctly identified a large fraction of contigs derived from ARG-plasmids (Recall_(contig)_=0.941), and considerably outperformed all individual classifiers. Thus, we believe that plasmidEC will be especially useful for plasmidome research that focuses on antibiotic resistance. Notably, all binary classifiers presented higher recall_(contig)_ for classifying contigs from ARG plasmids than from non-ARG plasmids, suggesting that these sequences might be overrepresented in reference databases which are directly or indirectly used by all tools.

When comparing the performance of the tools using the entire benchmark dataset, we found that plasmidEC and PlaScope performed very similarly in terms of F1-Score_(contig)_. However, plasmidEC showed a higher recall_(contig)_ but used more computational resources and took a longer time to complete the predictions. Reference-based methods, like PlaScope, are expected to perform well for species like *E. coli* which are abundant in public databases [39]. Supporting this hypothesis, a recent study by Shaw et al. [40] discovered very few novel plasmid sequences in a dataset that included more than 2,000 plasmids from *Enterobacteriaceae* isolates. PlaScope was built around Centrifuge [41], a metagenomic classifier to predict the origin of contigs based on custom databases. Recently, it was also shown that the usage of Kraken [42], another metagenomic classifier using customised databases, outperformed other binary classifiers in *Klebsiella pneumoniae [41,43]*. It would be interesting to explore how tools perform at classifying contigs from species with a limited number of complete genomes in databases. We speculate that in those cases, plasmidEC, which combines tools with diverse computational approaches, could improve predictions to a larger extent.

PlasmidEC could be further optimised by (i) multithreading the predictions of the individual tools, which would reduce the computational time to generate the results, (ii) including the possibility to predict the origin of contigs from other species, as long as those are supported by the binary classifiers, and (iii) improving its accuracy by using weighted votes, where a high confidence prediction will contribute more to the final result than a low confidence prediction.

We integrated plasmidEC (and PlaScope) with gplas2 to reconstruct individual *E. coli* plasmids. We then compared the performance of gplas2 combined with those classifiers against MOB-suite. Interestingly, the most pronounced differences in performance were observed when reconstructing ARG-plasmids. Although combined completeness_(bp)_ values indicated that the three tools identified similar fractions of ARG-plasmids, MOB-suite more frequently fragmented ARG-plasmids into multiple bins, yielding low completeness_(bp)_ and F1-Score_(bp)_. In contrast, gplas2 (either with plasmidEC or PlaScope) was more successful at binning together contigs into individual plasmid predictions, thus achieving considerably higher values for the aforementioned metrics. Accuracy_(bp)_ values for all tools were very similar, indicating a similar degree of chimeric predictions. Interestingly, both gplas2 methods performed similarly to MOB-suite when reconstructing plasmids that carry ESBL genes, which suggests that these plasmids might be overrepresented in the database used by MOB-suite to make predictions.

We recently described that ARG plasmids from *E. coli* are particularly difficult to reconstruct from short-read data [18], and we suggested that the modular nature of these plasmids could complicate their reconstruction using strict reference-based methods, such as MOB-suite. The results we obtained here seem to confirm this hypothesis. Additionally, we improved the reconstruction of ARG-plasmids by using coverage and node connectivity information. Yet, our study also proves that enriching the assembly graph with accurate information on the origin of contigs (plasmid/chromosome) is equally important. A previous version of gplas, which used mlplasmids as a binary classifier, performed significantly worse at predicting ARG-plasmids in *E. coli [19]*. Moreover, using a simpler graph-based approach that mainly relies on coverage differences to identify plasmids is also insufficient. This approach, applied by plasmidSPAdes, frequently leads to the inclusion of chromosomal contamination [18,19], due to the low copy number that ARG-plasmids often exhibit.

We envision that gplas2 could be combined with different binary classification tools to obtain accurate *de novo* plasmid reconstructions for multiple bacterial species. This means that gplas2 could, in theory, also be applied to the reconstruction of plasmids in metagenomic samples. However, since a greater number of plasmid-predicted unitigs is expected on metagenomes, the construction of plasmid walks will probably require parallelization in order to keep the computation time within practical limits.

Although our method constitutes a considerable improvement of the reconstruction of ARG-plasmids, some limitations should be noted. First, gplas2 does not include insertion sequences (and other repeated elements) into plasmid predictions. This facilitates the process of finding plasmid walks with homogeneous coverages and simplifies the resulting plasmidome network. However, insertion sequences play an important role in the structure and genomic plasticity of plasmids [44], and they are frequently involved in the mobility of ARGs [9,45,46]. Additionally, the localization of these MGEs can influence the expression levels of ARGs [47,48], thereby impacting the resulting resistance phenotypes. Consequently, including IS elements would certainly improve the completeness and relevance of plasmid predictions. Some graph-based plasmid reconstruction methods, like HyAsP [49], include repeated elements into predictions. This tool also constructs plasmid walks, and uses coverage information to predict IS copy numbers, thus allowing the same IS to be present in multiple replicons. In the gplas algorithm, considering repeated elements during the construction of the plasmid walks would lead to more entangled plasmidome networks and would complicate the subsequent partitioning step. As an alternative, we could envision adding labels to unitigs after the binning step, and then implementing a label propagation algorithm on the original assembly graph to determine to which bin the different IS elements belong. A similar approach is implemented by the tool GraphBin2 [50], which refines binning results of metagenomics samples. A second disadvantage of our method is the formation of chimeras, which are bins composed of nodes from distinct replicons. As previously mentioned, accurate identification of plasmid derived nodes reduces the number of chromosome-plasmid chimeras. However, preventing the formation of plasmid-plasmid chimeras is more challenging, especially for isolates carrying multiple large plasmids with similar copy numbers. Separating these chimeras could be possible with the use of a plasmid-backbone reference database.

To conclude, in this work we presented a new plasmidome prediction tool, named plasmidEC, and optimised gplas to accurately bin predicted plasmid sequences. Compared to existing binary classifiers, plasmidEC achieves increased recall_(contig)_, especially for contigs that derive from ARG plasmids. The integration of plasmidEC with gplas2 substantially improved the reconstruction of ARG plasmids in *E. coli*. Our method exceeded the binning capacity of the reference-based method MOB-suite, while retaining similar accuracy_(bp)_ values. The presented approach constitutes the best alternative to accurately predict and reconstruct ARG plasmids *de novo* in the absence of long-read data.

## Authors contributions

Conceptualization, J.A.P., A.C.S, S.A.A.; methodology, J.A.P., L.V, J.J.K.,S.A.A.; validation and formal analysis, J.A.P., L.V, J.J.K.; resources, supervision and project administration, A.C.S, S.A.A., R.J.L.W, N.L.P; data curation, J.A.P., L.V, .; writing—original draft preparation, J.A.P.; writing—review and editing, J.A.P., A.C.S., N.L.P.; visualisation, J.A.P., L.V. All authors have read and agreed to the published version of the manuscript.

