## Supplementary Material for "PlasmidEC and gplas2: An optimised short-read approach to predict and reconstruct antibiotic resistance plasmids in *Escherichia coli*"

### Supplementary Data

Tables including data from isolates used for the benchmark of plasmid reconstruction tools can be downloaded from: https://doi.org/10.5281/zenodo.7926472

### Supplementary Results

Fractions of large plasmids can be found on nodes smaller than 1kb

We used nodes larger than 1kb as an input for gplas2 and MOB-suite. When aligning all these nodes to their corresponding complete genomes, we discovered that a considerable fraction of some plasmids were missing (Supplementary Figure S7). The recovered fraction for small plasmids (median=1, IQR=0.98 - 1) was generally higher than for large plasmids (median=0.96, IQR= 0.91 - 0.99 ).

After including nodes with sizes ranging from 500 bp to 1 kb, we observed an increase in the recovered fraction to a total of 201 plasmids. This increase mainly occured in large plasmids (n=194, 71.9%) and rarely in small plasmids (n=7, 3.28%) (Supplementary Figure S7). However, the relative increase in recovered fraction within large plasmids was minimal (median=0.98, IQR=0.94 - 1).

Despite the inclusion of smaller contigs, the recovered fraction remained below 0.9 for 26 (9.62%) large plasmids. Additionally, a total of 2 small plasmids and 1 large plasmid were entirely missing after assembly (recovered fraction = 0).

In order to determine if the missing plasmids or plasmid regions were successfully sequenced, we aligned Illumina reads back to their complete genomes and analyzed the sequencing coverage distribution. For simplification, we show results for plasmids that had a recovered fraction below 0.8 (n=11). Most of these isolates contained multiple plasmids, presenting an overall median of 6 plasmids. We found Illumina reads aligning to all bases that were missed from assembly (Supplementary Figure S8). Interestingly, in 5 of these plasmids (CP051632.1, CP055630.1, AP022225.1, AP022249.1, CP054283.1) the median sequencing coverage was lower than the median coverage of the chromosome. In the remaining 6 cases, however, there was no apparent correlation between the coverage in unassembled regions and the chromosome median coverage (Supplementary Figure S9).

Given that recovered fractions for large plasmids increased when including smaller contigs, we re-run all plasmid prediction tools including these contigs as input. In contrast to expectation, the overall performance of the plasmid reconstruction tools did not improve and remained almost identical for every metric (Supplementary Figure S10).

### Supplementary Tables

**Supplementary Table S1.** Performance metrics of binary classifiers and plasmidEC combinations evaluated for complete benchmarking dataset. Predictions were categorised into: True Positives (TP, prediction = plasmid, class = plasmid), True Negatives (TN, prediction = chromosome, class = chromosome), False Positives (FP, prediction = plasmid, class = chromosome) and False Negatives (FN, prediction = chromosome, class = plasmid).

| Software | TP | TN | FP | FN | Precision | Recall | F1-Score |
|---|---|---|---|---|---|---|---|
| mlplasmids | 1,297 | 12,449 | 418 | 577 | 0.756 | 0.692 | 0.722 |
| PlaScope | 1,629 | 12,755 | 117 | 245 | 0.932 | 0.869 | 0.900 |
| Platon | 1,509 | 12,748 | 122 | 365 | 0.925 | 0.805 | 0.861 |
| RFPlasmid | 1,523 | 12,452 | 420 | 351 | 0.783 | 0.812 | 0.798 |
| mlplasmids/PlaScope/RFPlasmid | 1,588 | 12,689 | 183 | 286 | 0.896 | 0.847 | 0.871 |
| mlplasmids/Platon/PlaScope | 1,588 | 12,769 | 103 | 286 | 0.939 | 0.847 | 0.890 |
| mlplasmids/Platon/RFPlasmid | 1,536 | 12,694 | 178 | 338 | 0.896 | 0.819 | 0.560 |
| Platon/PlaScope/RFPlasmid | 1,658 | 12,736 | 136 | 216 | 0.924 | 0.884 | 0.904 |

**Supplementary Table S2.** Performance metrics of binary classifiers and plasmidEC combinations evaluated for ARG-plasmids. Predictions were categorised into: True Positives (TP, prediction = plasmid, class = plasmid) and False Negatives (FN, prediction = chromosome, class = plasmid).

| Software | TP | FN | Recall |
|---|---|---|---|
| mlplasmids | 622 | 238 | 0.723 |
| PlaScope | 760 | 100 | 0.883 |
| Platon | 738 | 122 | 0.858 |
| RFPlasmid | 731 | 129 | 0.85 |
| mlplasmids/PlaScope/RFPlasmid | 760 | 100 | 0.883 |
| mlplasmids/Platon/PlaScope | 768 | 92 | 0.893 |
| mlplasmids/Platon/RFPlasmid | 753 | 107 | 0.875 |
| Platon/PlaScope/RFPlasmid | 809 | 51 | 0.941 |

### Supplementary Figures

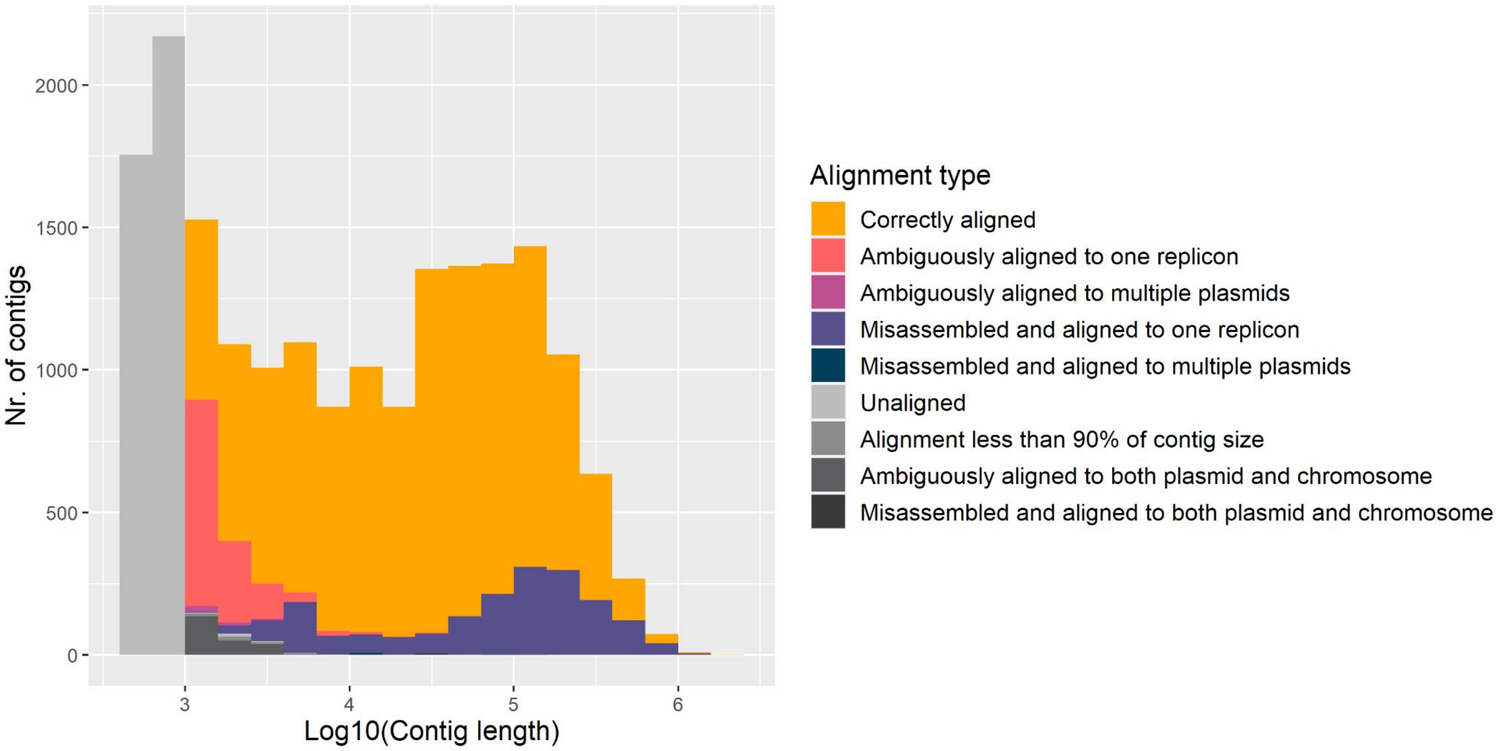


**Supplementary Figure S1.** Alignment types of all contigs in the dataset by contig length. Included contigs are shown in colour, excluded contigs in greyscale.

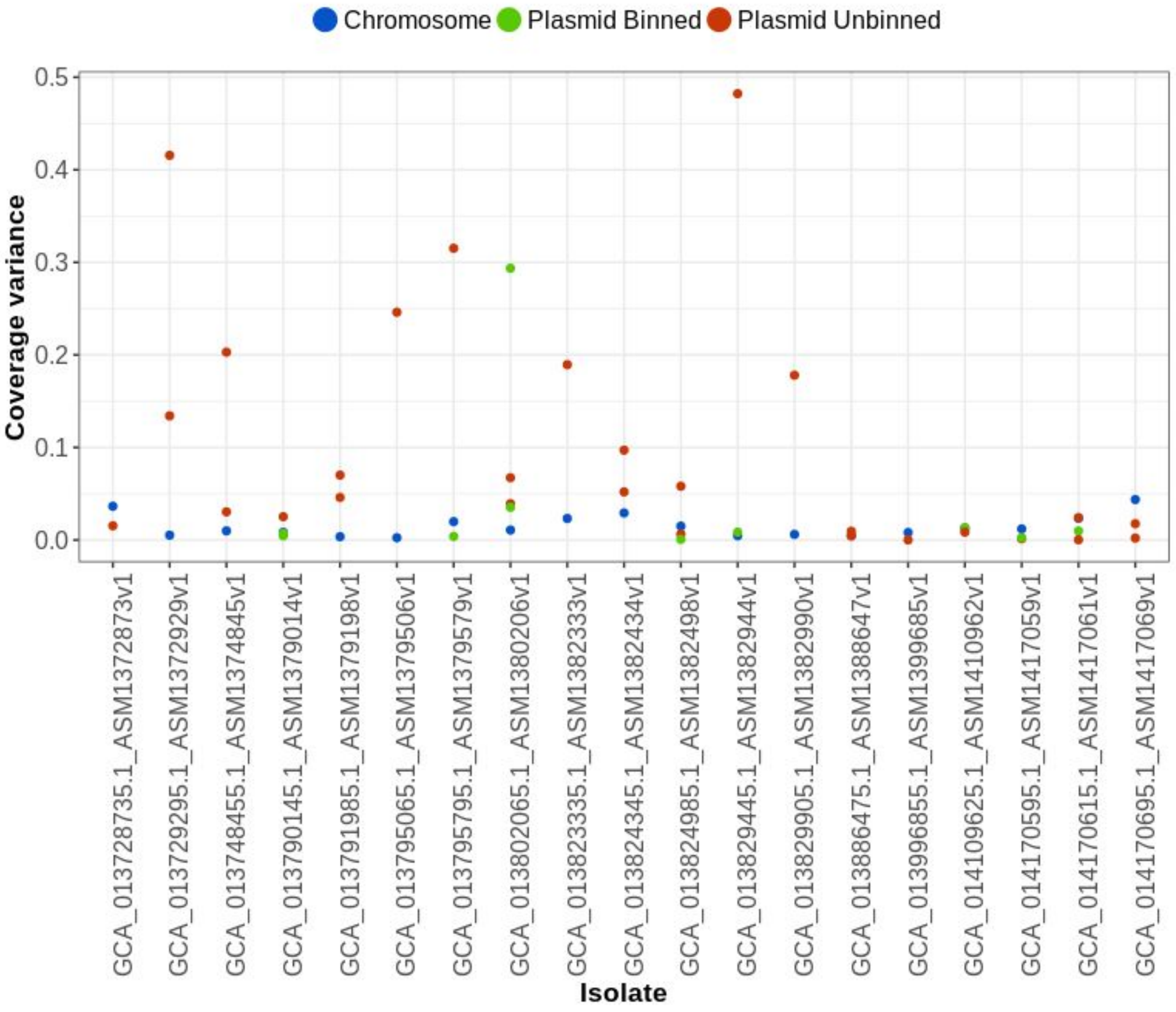


**Supplementary Figure S2.** Contig coverage variance for all replicons carried by isolates that contained unbinned nodes after gplas_plasmidEC prediction.

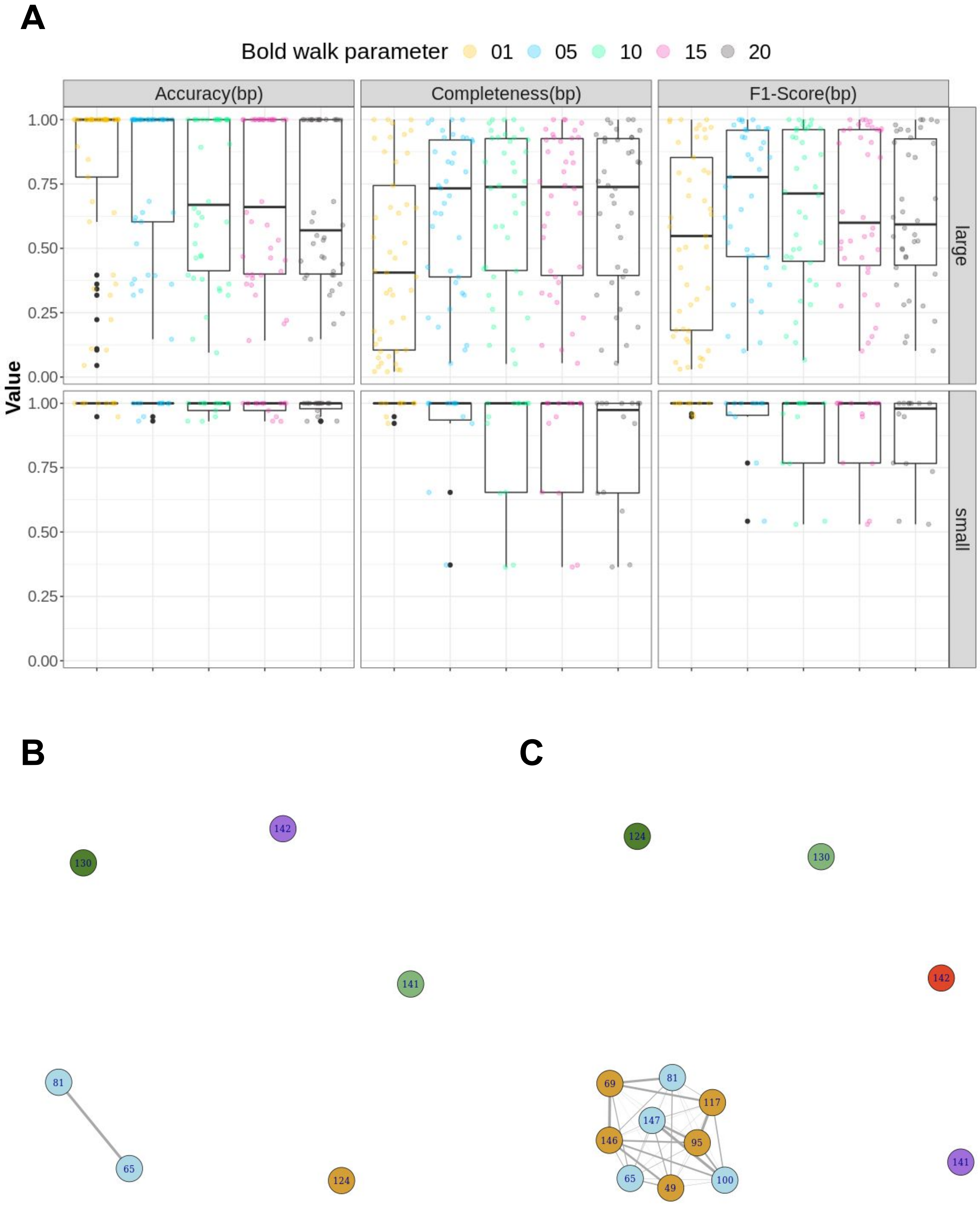


**Supplementary Figure S3.** A) Completeness(bp), Accuracy(bp) and F1-score(bp) values for plasmid predictions derived from isolates in which unbinned unitigs were predicted by gplas_plasmidEC (n=15). gplas_plasmidEC was run allowing different coverage variances in the bold mode. B) Plasmidome network obtained after running gplas2 without the bold parameter on isolate GCA_013823335.1_ASM1382333v1. Circles represent unitigs predicted as plasmid by plasmidEC and binned by gplas. Different colours correspond to different individual predicted plasmids. C) Plasmidome network obtained after running gplas2 with bold parameter of 5, on the same isolate.

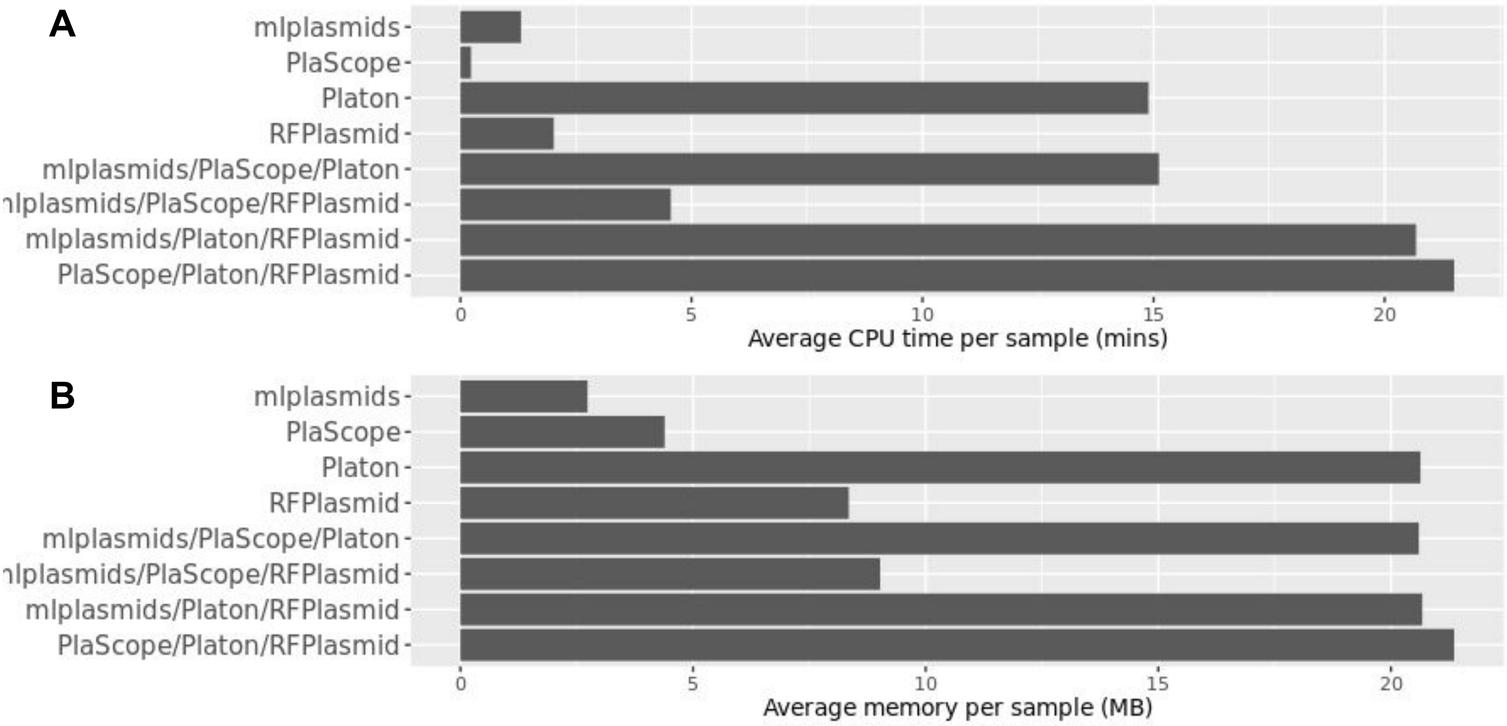


**Supplementary Figure S4.** Average computational resources used per sample: CPU time in minutes (A) and memory in Mb (B). Softwares were run on the full benchmarking dataset (n = 214).

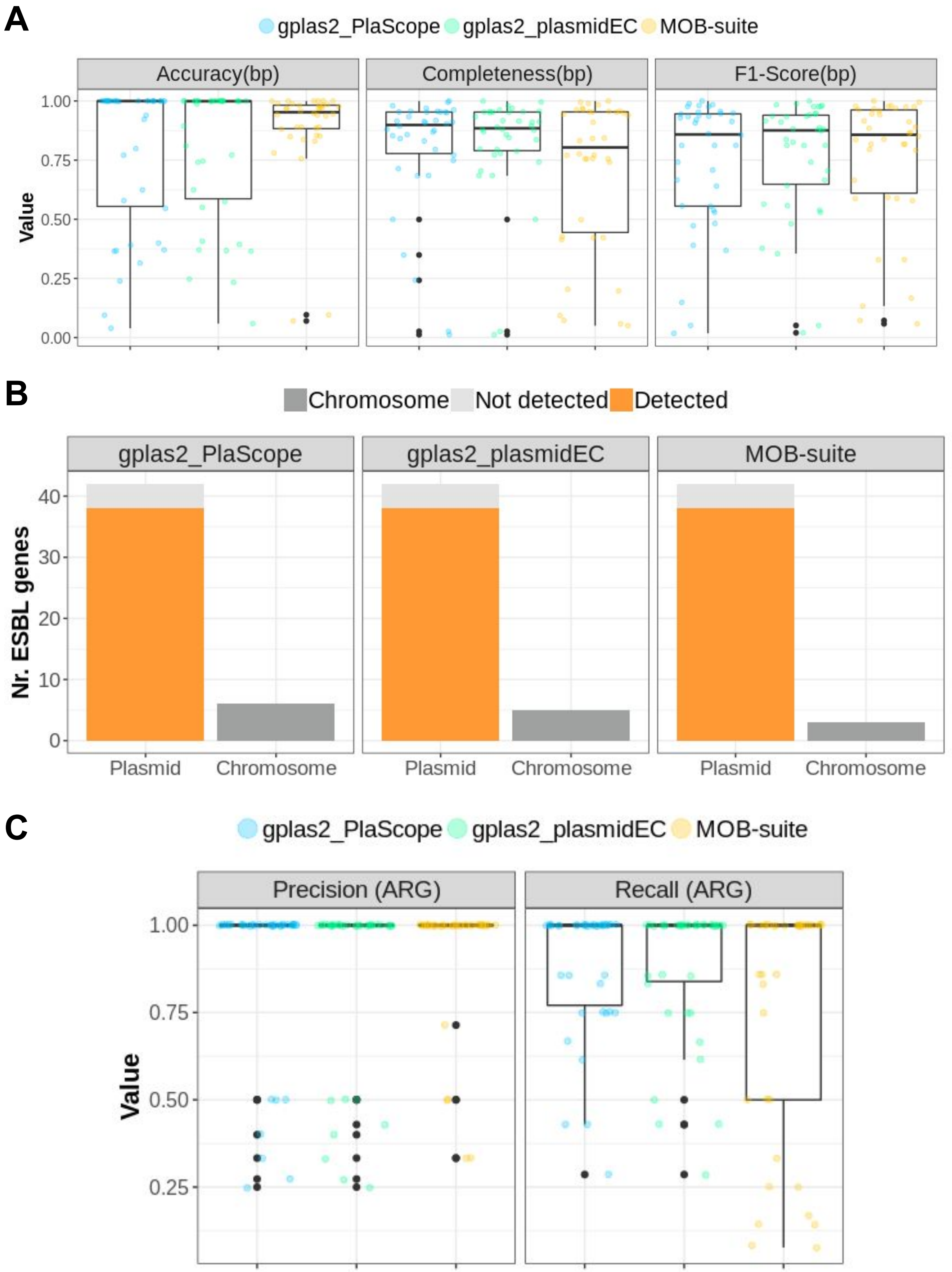


**Supplementary Figure S5.** A) Completeness(bp), Accuracy(bp), F1-score(bp) values for predictions carrying plasmid-derived ESBL genes (n=42). B) Absolute count of plasmid-derived ESBL genes included (detected) and missed (Not detected) in plasmid predictions. Absolute count of chromosome-derived ESBLs contaminating plasmid predictions are also depicted. C) Recall(ARG) and Precision(ARG) values for ESBL-carrying plasmids.

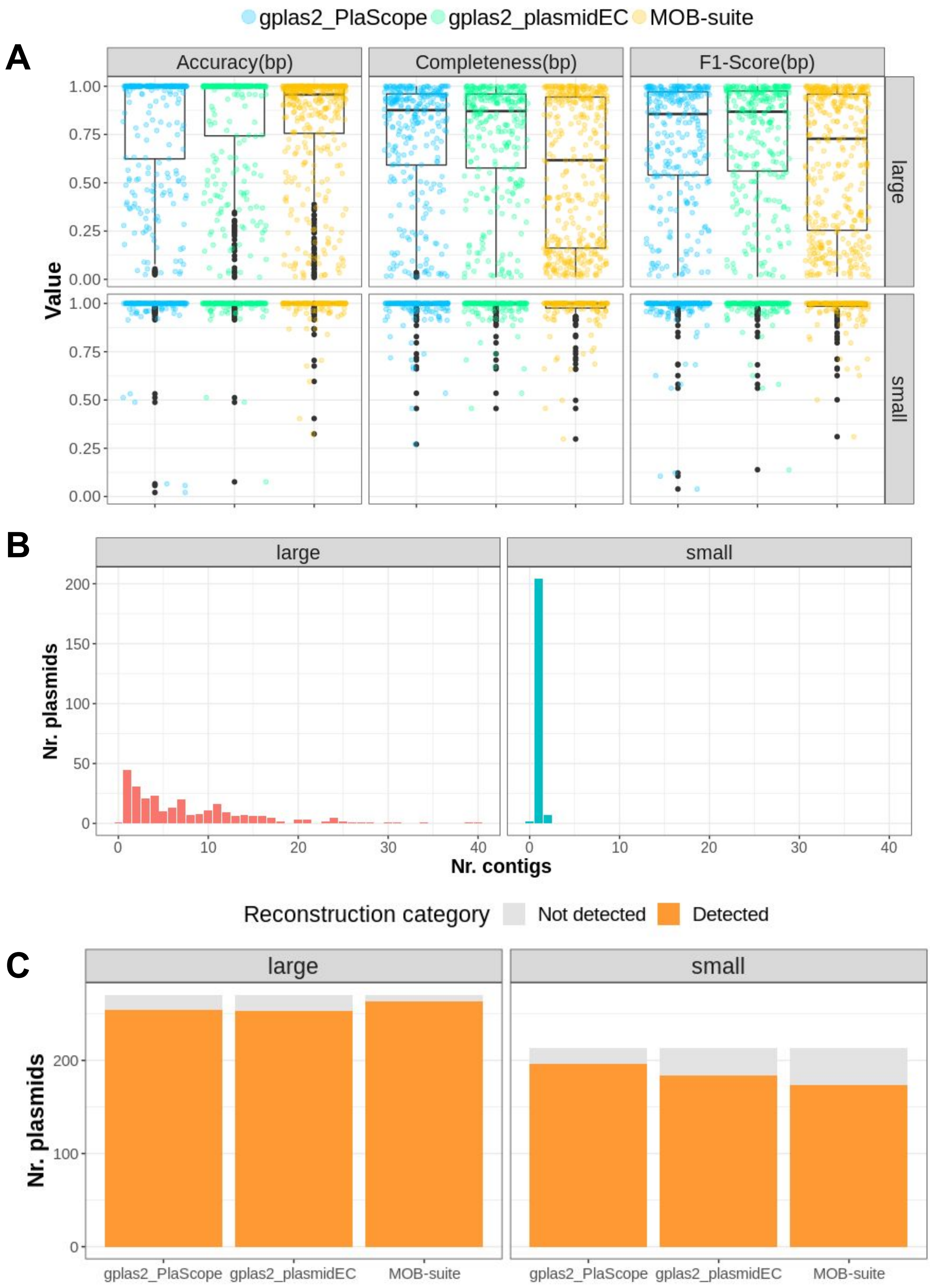


**Supplementary Figure S6.** A) Completeness(bp), Accuracy(bp), F1-score(bp) values for plasmid predictions according to size category (definition: small plasmids <18 kp, large plasmids >=18kb). B) Histogram showing the number of contigs that compose each reference plasmids when these are assembled from short reads only. C) Absolute count of plasmids detected and undetected by each of the tools according to plasmid size. A reference plasmid was labelled as detected when at least one of its contigs was included into the predictions.

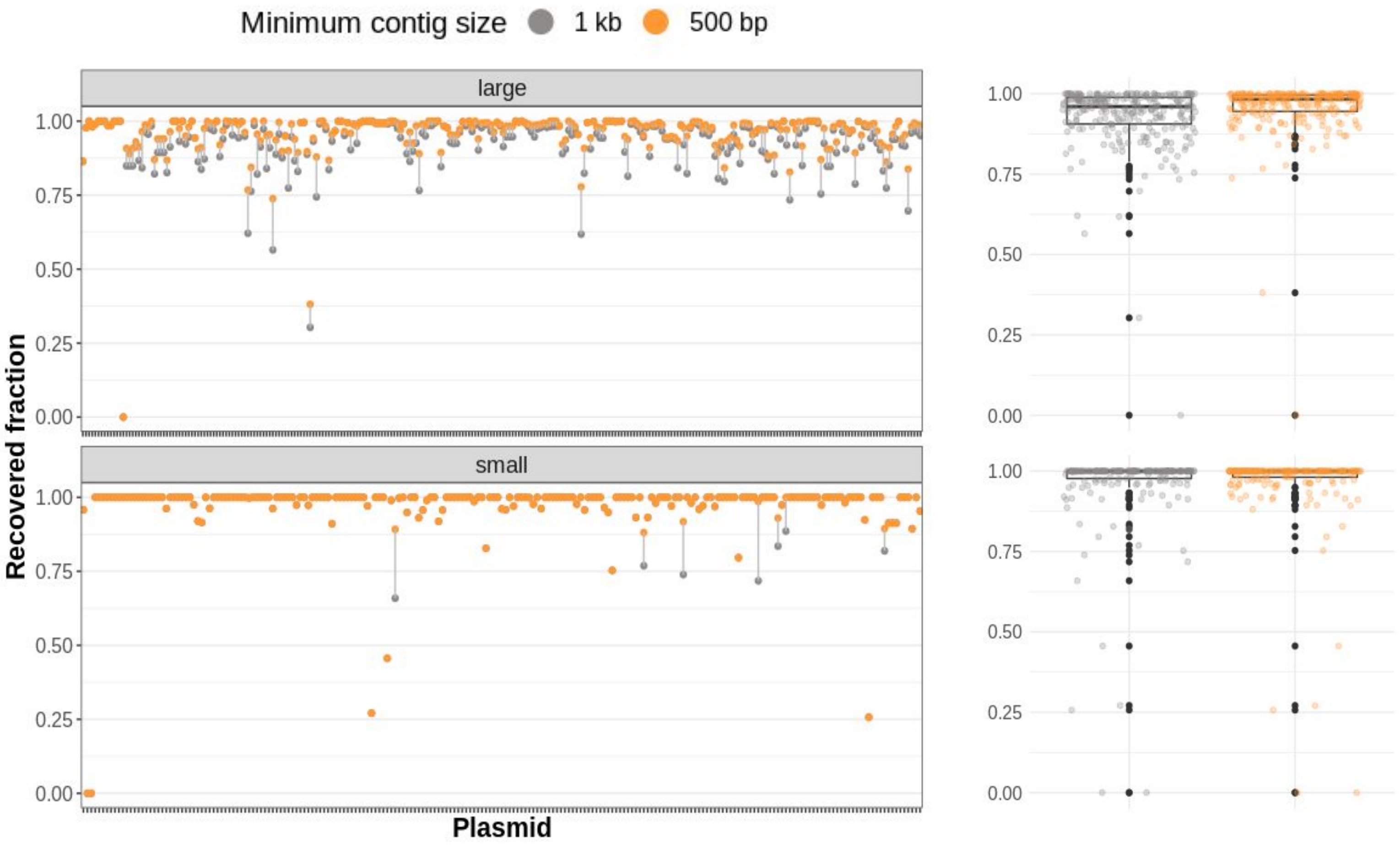


**Supplementary Figure S7.** Recovered fraction of plasmids after aligning all contigs larger than 500 bp (orange) or larger than 1 kb (grey) to the complete genomes.

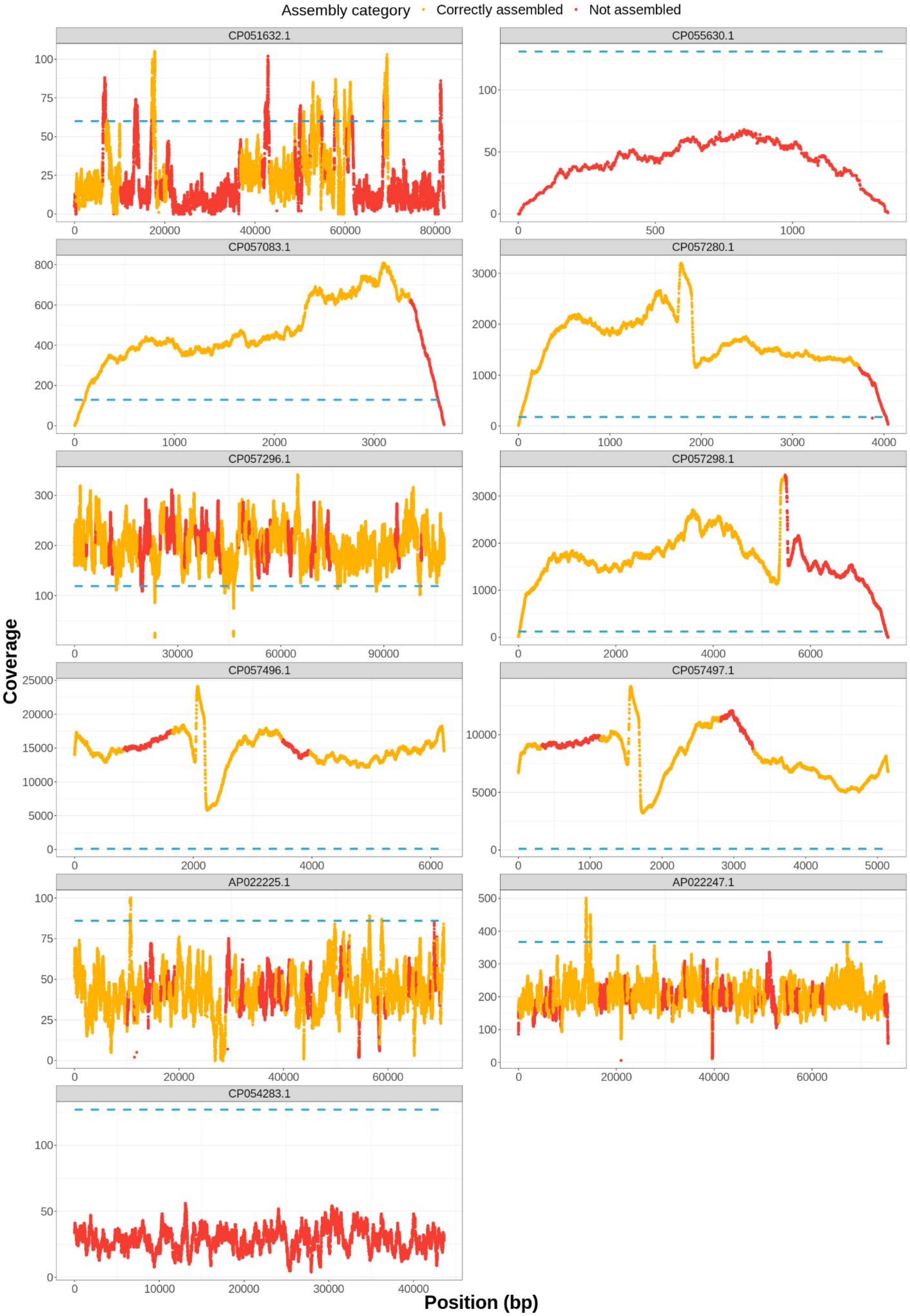


**Supplementary Figure S8.** Read coverage per replicon position for plasmids that had a recovered fraction equal or smaller than 0.8. Regions that were not assembled in contigs larger than 500 bp are shown in red. Blue dotted line represents the median coverage for the chromosome.

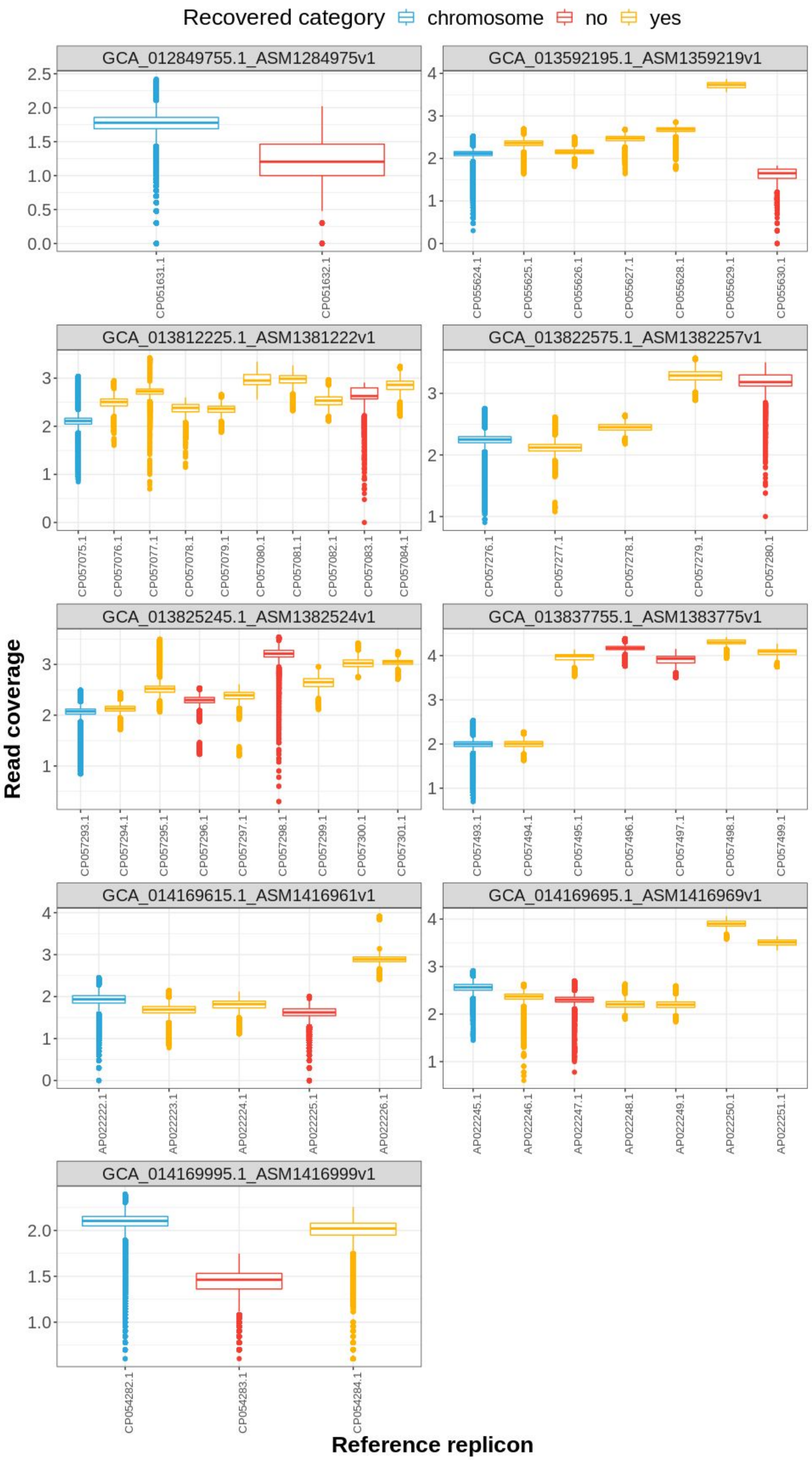


**Supplementary Figure S9.** Read coverage distribution for all replicons carried by isolates that had at least one plasmid with a recovered fraction <0.8 (n=11).

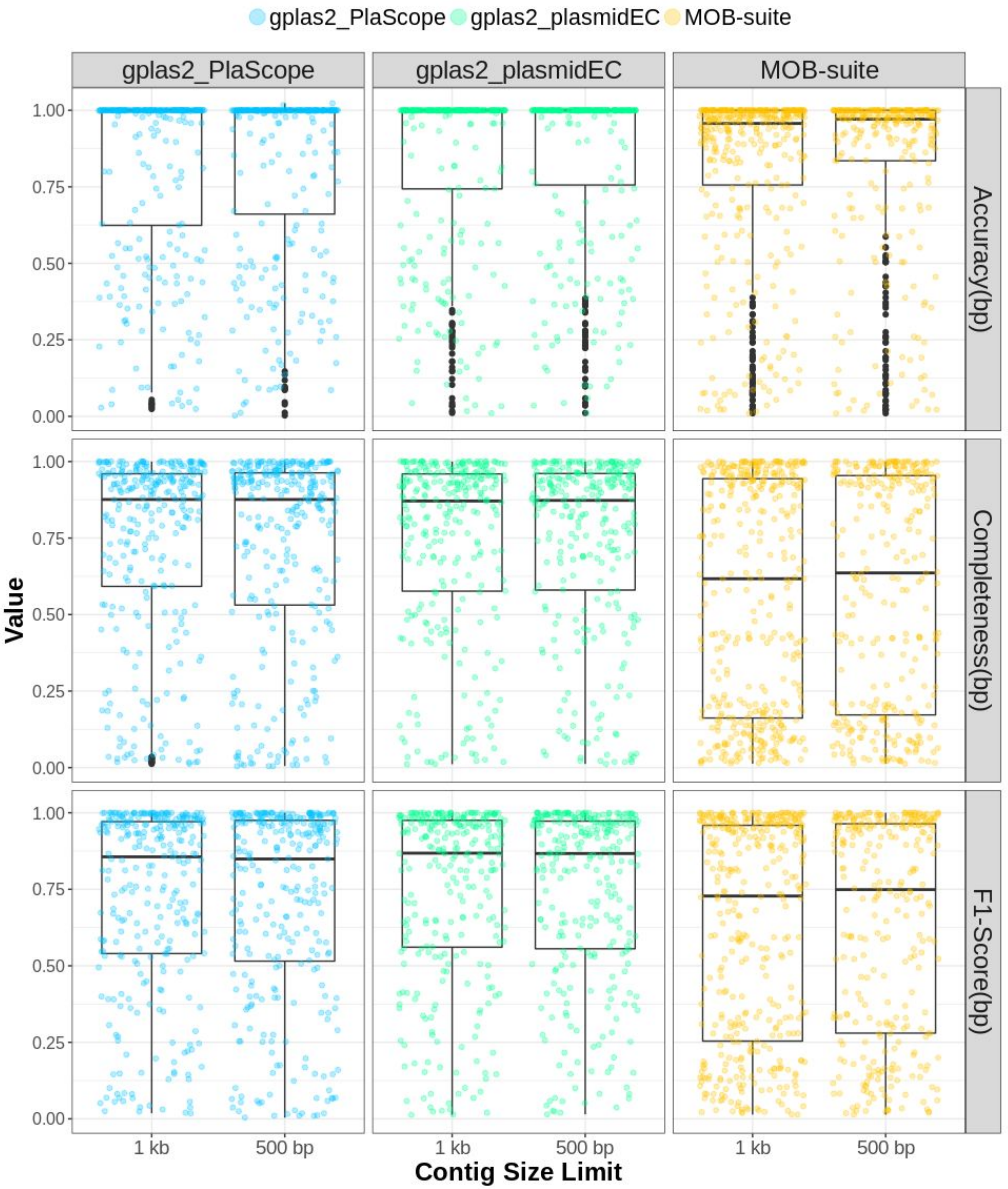


**Supplementary Figure S10.** Distribution of Accuracy(bp), Completeness(bp) and F1-Score(bp) values for plasmid predictions obtained using as input contigs of sizes larger than 500 bp or larger than 1 kb. Only large plasmids (n=270) were included in this analysis.
